# Hypoxemia is an Independent Risk Factor for Perioperative Thrombosis at High Altitude: A Retrospective Study of 11,084 Patients

**DOI:** 10.64898/2026.11.04.764430

**Authors:** Jin Yeyuan, Wu YiFan, Ma SuXin, Peng ChunChao, Li Xiang, He Zongzhao

**Affiliations:** Department of anesthesiology, Qinghai provincal people’s hospital, Xi’ning, China; Qinghai University

**Keywords:** High altitude, Perioperative period, Hypoxemia, Venous thromboembolism, Logistic regression, Risk factors

## Abstract

**Objective:** Perioperative thrombosis occurs at a significantly higher rate at high altitudes compared to low altitudes, severely threatening the safety of surgical patients. This study aims to determine the independent risk factors for perioperative thrombosis in high-altitude environments, providing a scientific basis for early clinical screening and targeted intervention.

**Methods:** Clinical data from 11,084 surgical patients were retrospectively analyzed. Patients were categorized into thrombosis and non-thrombosis groups based on the occurrence of postoperative thrombosis. Baseline characteristics, operative variables, and laboratory indicators were collected, encompassing 21 potential risk factors including arterial partial pressure of oxygen (PaO_2_). Univariate analysis and multivariate logistic regression (forward stepwise likelihood ratio method) were performed using SPSS 27.0 to identify independent risk factors for perioperative thrombosis. The predictive performance of the model was evaluated using receiver operating characteristic (ROC) curves, decision curve analysis (DCA), calibration plots, and nomograms.

**Results:** Of the 11,084 patients, 1,631 (14.7%) developed thrombosis. Univariate analysis showed that varicose veins, hemoglobin levels, Preoperative hospital stay, body mass index (BMI), D-dimer, partial pressure of oxygen, and interleukin-6 were significantly associated with thrombosis (*P* < 0.05). Multivariate logistic regression identified varicose veins, Preoperative hospital stay, elevated BMI, erythrocytosis, elevated D-dimer, hypoxemia, and elevated interleukin-6 as independent risk factors for thrombosis (*P* < 0.05). The predictive model demonstrated strong discrimination, with an area under the curve (*AUC*) of 0.862 (95% CI: 0.848–0.876), a sensitivity of 0.790, and a specificity of 0.773. Among the individual risk factors, hypoxemia exhibited the highest diagnostic performance, yielding an *AUC* of 0.732 and a sensitivity of 0.763.

**Conclusion:** The incidence of perioperative thrombosis at high altitudes is elevated, with hypoxemia serving as an independent predictor beyond traditional risk factors. Clinical management should prioritize correcting pre– and postoperative hypoxic states. Furthermore, implementing individualized prophylaxis that accounts for other risk factors such as multiple surgeries, advanced age, anemia, and coagulation-fibrinolysis system abnormalities is crucial to mitigating thrombotic risk.

## Introduction

Perioperative thrombosis represents a critical complication of surgical procedures; it not only prolongs hospital stays and escalates healthcare costs but can also culminate in fatal pulmonary embolism, severely threatening patient safety[1].

High-altitude environments are characterized by unique stressors, including hypoxia, hypobaric pressure, and aridity. Long-term high-altitude residents undergo adaptive physiological transformations driven by chronic hypoxic stimulation, such as compensatory erythrocytosis, hyperviscosity, pulmonary vasoconstriction, and endothelial dysfunction[2, 3]. Although these adaptations facilitate acclimatization to low-oxygen environments, they concurrently elevate thrombotic risk [4].

Consequently, we hypothesize that residential altitude and hypoxemia serve as independent risk factors for perioperative thrombosis in these regions. However, large-scale, multifactorial studies addressing this issue remain limited. Conventional preoperative risk assessment models widely utilized in clinical practice (e.g., the Caprini score) were primarily validated in sea-level cohorts and fail to incorporate altitude-specific variables. To address this knowledge gap, we conducted a large-sample retrospective analysis to identify the independent risk factors for perioperative thrombosis in a high-altitude population.

## Methods

### 1.1 Study Population and Data Sources

This retrospective study received ethical approval and exemption.This retrospective study included patients who underwent surgical procedures at Qinghai Provincial People’s Hospital (altitude ∼2,200 meters) between December 2024 and December 2025., The inclusion criteria were: (1) age ≥ 18 years; (2) receipt of surgical intervention during hospitalization; and (3) complete clinical data, including perioperative extremity vascular ultrasound and arterial blood gas analysis. Patients were excluded if they: (1) had a confirmed diagnosis of thrombosis prior to surgery; or (2) had severely incomplete clinical records. De-identified datasets were extracted from the electronic medical record system by institutional information department engineers. Because this study utilized a fully de-identified retrospective dataset, it was deemed exempt from Institutional Review Board (IRB) review. Ultimately, 11,084 patients were included and categorized into either a thrombosis group (n = 1,631) or a non-thrombosis group (n = 9,453) based on postoperative color Doppler ultrasound confirmation during their hospital stay.

### 1.2 Outcome Definitions

Perioperative thrombosis was defined as any arterial or venous thrombus detected via vascular color Doppler ultrasonography during hospitalization. This encompassed deep vein thrombosis, intermuscular vein thrombosis, intramuscular venous plexus thrombosis, and arterial thrombosis. Sonographic diagnosis of a thrombus required the presence of the following three key imaging features: (1) incomplete vein compressibility under transducer pressure; (2) abnormal intraluminal hyperechoic structures; and (3) absence or marked attenuation of blood flow signals on color Doppler imaging at the suspected site.

### 1.3 Covariates for Risk Modeling

Baseline covariates comprised patient characteristics, laboratory parameters, and arterial blood gas (ABG) findings.

Patient characteristics included age, sex, body mass index (BMI, weight/height^2^), altitude(high-altitude(1,500–2,500 meters) and plateau (>2,500 meters)),surgical frequency (single vs. multiple surgeries, with the latter defined as ≥ 2 procedures during the index hospitalization; categorical), comorbidities (including cardiovascular, pulmonary, cerebrovascular, and renal diseases, diabetes, prior surgical history, and presence of multiple complications; categorical), history of varicose veins (yes vs. no), anesthesia modality (general, neuraxial, local, brachial plexus block, or non-intubated general anesthesia; categorical), and Preoperative hospital stay(LOS; continuous).

Laboratory parameters were obtained from blood samples collected within 24 hours before or after color Doppler ultrasound examination, including: hemoglobin levels (normal, anemia, or polycythemia), platelet count (normal, decreased, or increased), prothrombin time (PT; normal, shortened, or prolonged), prothrombin activity percentage (normal vs. decreased), international normalized ratio (INR; normal vs. shortened), fibrinogen (FIB; normal, decreased, or increased), fibrin degradation products (FDP; normal vs. increased), D-dimer (normal vs. increased), procalcitonin (PCT; normal vs. elevated), and interleukin-6 (IL-6; normal vs. elevated), all analyzed as categorical variables.

ABG variables included partial pressure of carbon dioxide (PaCO_2_; normal, decreased, or increased; categorical) and partial pressure of oxygen (PaO_2_; normal vs. hypoxemia; categorical). Hypoxemia was defined as PaO_2_ < 60 mmHg based on sea-level standards. This unified threshold was applied because clinical blood gas analyzers in this high-altitude setting are typically not corrected for altitude.

### 1.4 Statistical Analysis

Statistical analyses were performed using SPSS version 27.0. Categorical variables are expressed as frequencies and percentages [n (%)] and compared between groups using the chi-square (χ^2^) test. The normality of continuous variables was assessed using the Kolmogorov-Smirnov test. Normally distributed data are presented as means ± standard deviations (mean ± SD) and analyzed via the independent-samples t-test. Non-normally distributed data are expressed as medians with interquartile ranges [medians (25th, 75th percentiles)] and compared using the Mann-Whitney U test. Statistical significance was defined as P < 0.05.

## Results

Demographic and clinical parameters were compared between patients with and without thrombosis. Categorical data (including surgical frequency, comorbidities, history of varicose veins, and sex) were evaluated using the chi-square (χ^2^) test. Continuous variables (including total length of hospital stay, age, and BMI) exhibited non-normal distributions based on the Kolmogorov-Smirnov test and were thus analyzed using the Mann-Whitney U test. Univariate analysis revealed that the prevalences or levels of varicose veins, hemoglobin, Preoperative hospital stay, BMI, D-dimer, partial pressure of oxygen (PaO_2_), and interleukin-6 (IL-6) differed significantly between the two groups (all *P* < 0.05). No statistically significant differences were observed for the remaining parameters (*P* > 0.05). (Table 1)

**Table 1.**
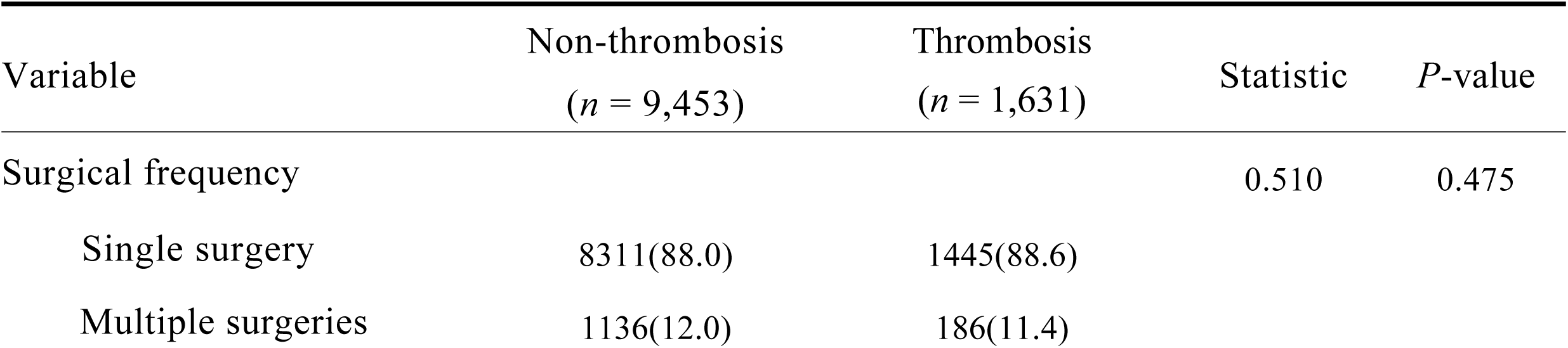
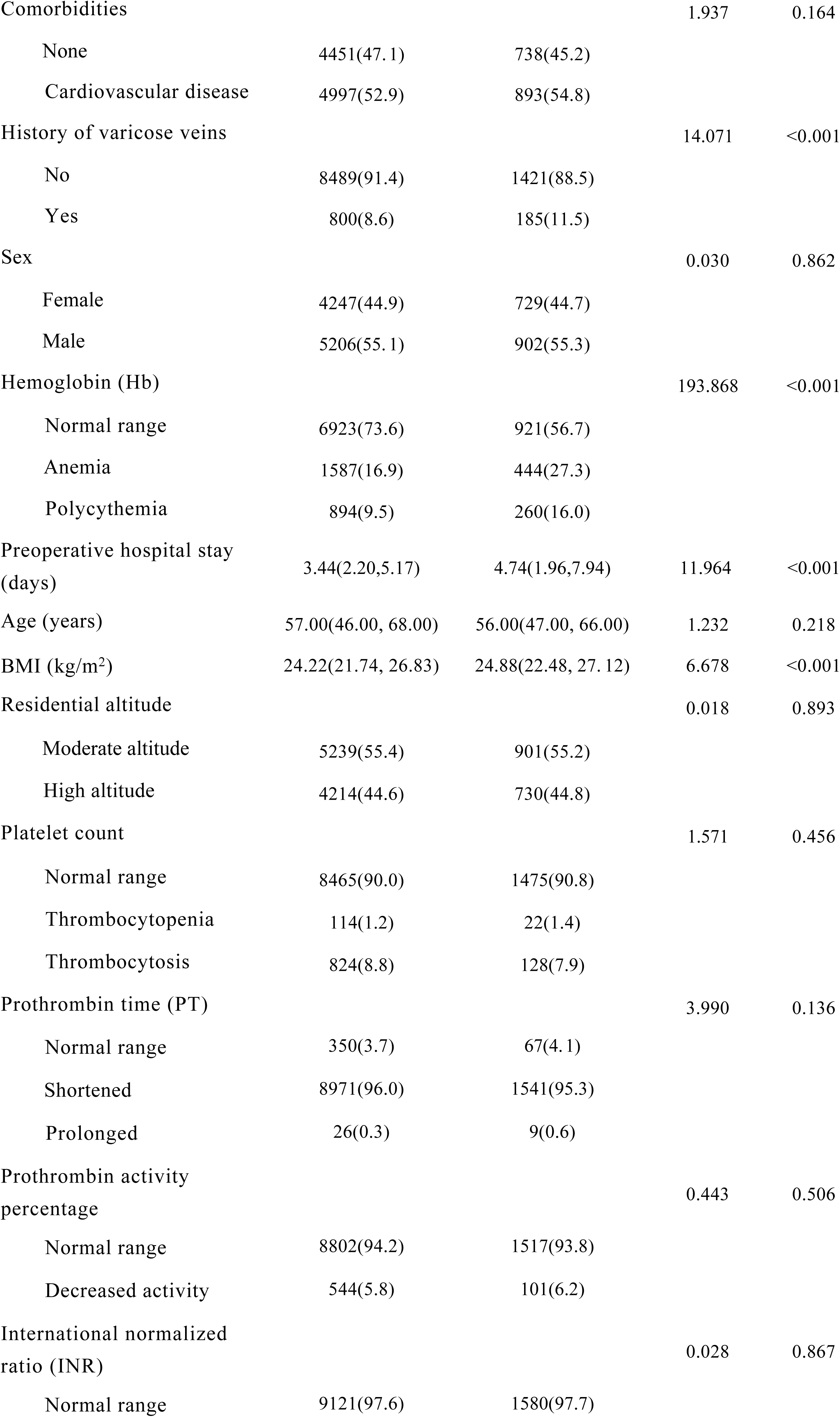
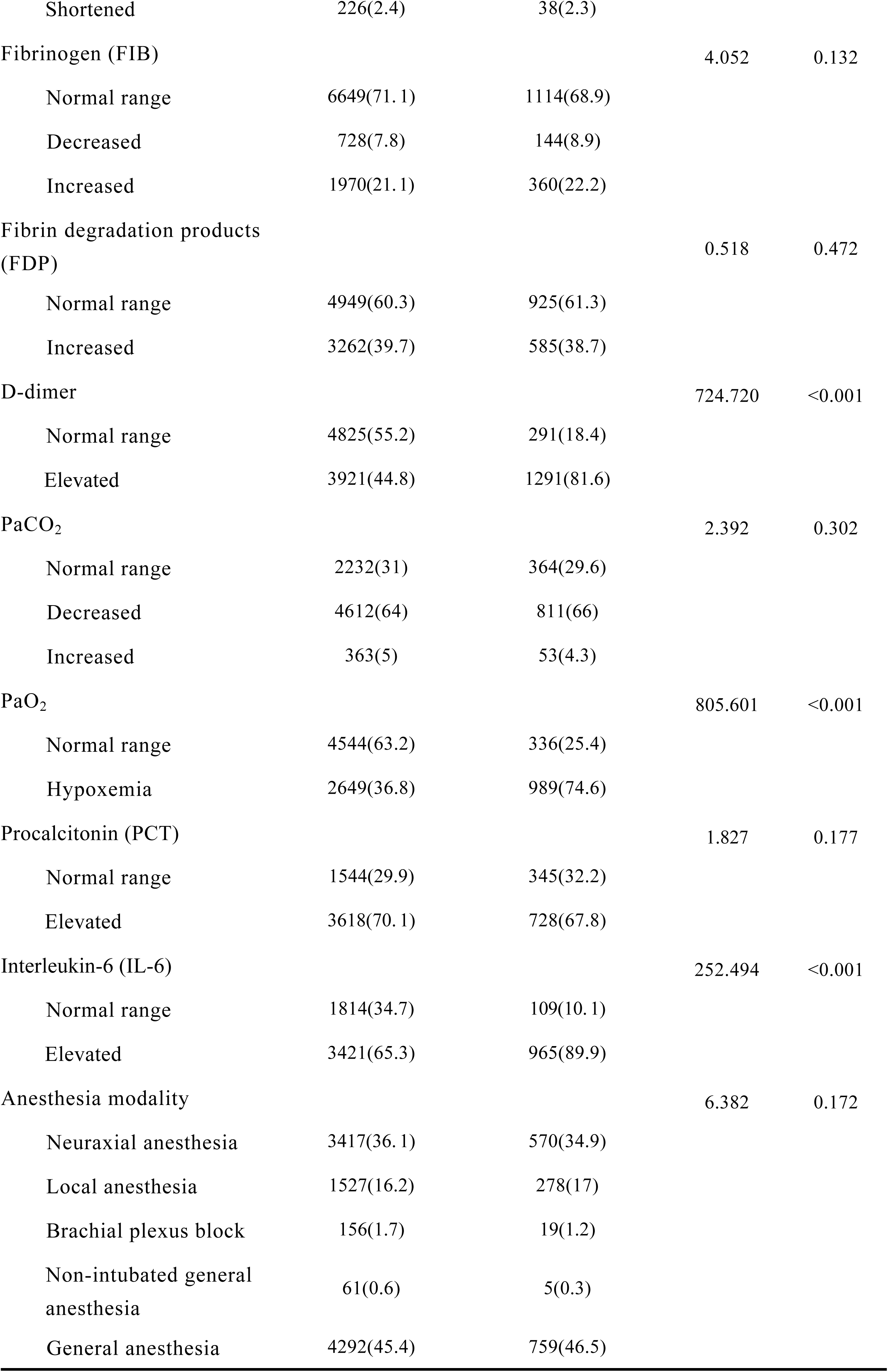
Univariate analysis of potential risk factors for perioperative thrombosis.

| Variable | Non-thrombosis<br>(*n* = 9,453) | Thrombosis<br>(*n* = 1,631) | Statistic | *P*-value |
|---|---|---|---|---|
| Surgical frequency |  |  | 0.510 | 0.475 |
| Single surgery | 8311(88.0) | 1445(88.6) |  |  |
| Multiple surgeries | 1136(12.0) | 186(11.4) |  |  |
| Comorbidities |  |  | 1.937 | 0.164 |
| None | 4451(47. 1) | 738(45.2) |  |  |
| Cardiovascular disease | 4997(52.9) | 893(54.8) |  |  |
| History of varicose veins |  |  | 14.071 | <0.001 |
| No | 8489(91.4) | 1421(88.5) |  |  |
| Yes | 800(8.6) | 185(11.5) |  |  |
| Sex |  |  | 0.030 | 0.862 |
| Female | 4247(44.9) | 729(44.7) |  |  |
| Male | 5206(55. 1) | 902(55.3) |  |  |
| Hemoglobin (Hb) |  |  | 193.868 | <0.001 |
| Normal range | 6923(73.6) | 921(56.7) |  |  |
| Anemia | 1587(16.9) | 444(27.3) |  |  |
| Polycythemia | 894(9.5) | 260(16.0) |  |  |
| Preoperative hospital stay (days) | 3.44(2.20,5.17) | 4.74(1.96,7.94) | 11.964 | <0.001 |
| Age (years) | 57.00(46.00, 68.00) | 56.00(47.00, 66.00) | 1.232 | 0.218 |
| BMI (kg/m<sup>2</sup>) | 24.22(21.74, 26.83) | 24.88(22.48, 27. 12) | 6.678 | <0.001 |
| Residential altitude |  |  | 0.018 | 0.893 |
| Moderate altitude | 5239(55.4) | 901(55.2) |  |  |
| High altitude | 4214(44.6) | 730(44.8) |  |  |
| Platelet count |  |  | 1.571 | 0.456 |
| Normal range | 8465(90.0) | 1475(90.8) |  |  |
| Thrombocytopenia | 114(1.2) | 22(1.4) |  |  |
| Thrombocytosis | 824(8.8) | 128(7.9) |  |  |
| Prothrombin time (PT) |  |  | 3.990 | 0.136 |
| Normal range | 350(3.7) | 67(4. 1) |  |  |
| Shortened | 8971(96.0) | 1541(95.3) |  |  |
| Prolonged | 26(0.3) | 9(0.6) |  |  |
| Prothrombin activity percentage |  |  | 0.443 | 0.506 |
| Normal range | 8802(94.2) | 1517(93.8) |  |  |
| Decreased activity | 544(5.8) | 101(6.2) |  |  |
| International normalized ratio (INR) |  |  | 0.028 | 0.867 |
| Normal range | 9121(97.6) | 1580(97.7) |  |  |
| Shortened | 226(2.4) | 38(2.3) |  |  |
| Fibrinogen (FIB) |  |  | 4.052 | 0.132 |
| Normal range | 6649(71. 1) | 1114(68.9) |  |  |
| Decreased | 728(7.8) | 144(8.9) |  |  |
| Increased | 1970(21. 1) | 360(22.2) |  |  |
| Fibrin degradation products (FDP) |  |  | 0.518 | 0.472 |
| Normal range | 4949(60.3) | 925(61.3) |  |  |
| Increased | 3262(39.7) | 585(38.7) |  |  |
| D-dimer |  |  | 724.720 | <0.001 |
| Normal range | 4825(55.2) | 291(18.4) |  |  |
| Elevated | 3921(44.8) | 1291(81.6) |  |  |
| PaCO<sub>2</sub> |  |  | 2.392 | 0.302 |
| Normal range | 2232(31) | 364(29.6) |  |  |
| Decreased | 4612(64) | 811(66) |  |  |
| Increased | 363(5) | 53(4.3) |  |  |
| PaO<sub>2</sub> |  |  | 805.601 | <0.001 |
| Normal range | 4544(63.2) | 336(25.4) |  |  |
| Hypoxemia | 2649(36.8) | 989(74.6) |  |  |
| Procalcitonin (PCT) |  |  | 1.827 | 0.177 |
| Normal range | 1544(29.9) | 345(32.2) |  |  |
| Elevated | 3618(70. 1) | 728(67.8) |  |  |
| Interleukin-6 (IL-6) |  |  | 252.494 | <0.001 |
| Normal range | 1814(34.7) | 109(10. 1) |  |  |
| Elevated | 3421(65.3) | 965(89.9) |  |  |
| Anesthesia modality |  |  | 6.382 | 0.172 |
| Neuraxial anesthesia | 3417(36. 1) | 570(34.9) |  |  |
| Local anesthesia | 1527(16.2) | 278(17) |  |  |
| Brachial plexus block | 156(1.7) | 19(1.2) |  |  |
| Non-intubated general anesthesia | 61(0.6) | 5(0.3) |  |  |
| General anesthesia | 4292(45.4) | 759(46.5) |  |  |

To identify independent risk factors for perioperative thrombosis, a multivariable logistic regression model was constructed. The model utilized the development of thrombosis (thrombosis = 1, non-thrombosis = 0) as the dependent variable and the statistically significant parameters from the univariate analysis as independent variables. The regression results demonstrated that a history of varicose veins,Preoperative hospital stay, BMI, polycythemia, elevated D-dimer, hypoxemia, and elevated interleukin-6 (IL-6) were independent risk factors of perioperative thrombosis formation (all *P* < 0.05; Table 2).

**Table 2.** Multivariable logistic regression analysis of risk factors for thrombosis.

| Variable | *β* | SE | Wald | *P* | OR | 95% CI |
|---|---|---|---|---|---|---|
| History of varicose veins | 0.394 | 0.164 | 5.754 | 0.016 | 1.483 | 1.075-2.047 |
| Preoperative hospital stay(days) | 0.075 | 0.011 | 48.158 | <0.001 | 1.078 | 1.056-1.102 |
| BMI (kg/m<sup>2</sup>) | 0.069 | 0.012 | 33.280 | <0.001 | 1.072 | 1.047-1.097 |
| Hb (*Reference: Normal range*) |  |  | 26.913 | <0.001 |  |  |
| Anemia | 0.112 | 0.116 | 0.922 | 0.337 | 1.118 | 0.89-1.404 |
| Polycythemia | 0.740 | 0.143 | 26.912 | <0.001 | 2.097 | 1.585-2.773 |
| Elevated D-dimer | 1.825 | 0.125 | 211.771 | <0.001 | 6.203 | 4.852-7.932 |
| Hypoxemia | 1.981 | 0.102 | 377.710 | <0.001 | 7.249 | 5.936-8.852 |
| Elevated IL-6 | 1.086 | 0.149 | 53.111 | <0.001 | 2.964 | 2.213-3.969 |
| Constant | -6.982 | 0.373 | 351.004 | <0.001 | 0.001 |  |

Receiver operating characteristic (ROC) curves were constructed for the independent risk factors identified in the multivariable analysis to evaluate their respective diagnostic capacities for perioperative thrombosis. The area under the curve (*AUC*), along with the paired sensitivity and specificity, was determined for each variable: a history of varicose veins (*AUC* = 0.503; sensitivity: 0.102, specificity: 0.905),Preoperative hospital stay (*AUC* = 0.560; sensitivity: 0.359, specificity: 0.838), polycythemia (*AUC* = 0.534; sensitivity: 0.175, specificity: 0.893), hypoxemia (*AUC* = 0.732; sensitivity: 0.763, specificity: 0.700), elevated interleukin-6 (*AUC* = 0.631; sensitivity: 0.908, specificity: 0.354), elevated D-dimer (*AUC* = 0.697; sensitivity: 0.851, specificity: 0.542), and BMI (*AUC* = 0.558; sensitivity: 0.695, specificity: 0.434). Among all evaluated metrics, hypoxemia yielded the optimal diagnostic performance.

**Table 3.** Diagnostic performance of different indicators for predicting thrombosis.

| Variable | *AUC* | *P* | Sensitivity | Specificity | Youden index |
|---|---|---|---|---|---|
| History of varicose veins | 0.503 | 0.772 | 0.102 | 0.905 | 0.007 |
| Preoperative hospital stay(days) | 0.560 | <0.001 | 0.359 | 0.838 | 0.197 |
| Polycythemia | 0.534 | 0.004 | 0.175 | 0.893 | 0.068 |
| Hypoxemia | 0.732 | <0.001 | 0.763 | 0.700 | 0.463 |
| Elevated IL-6 | 0.631 | <0.001 | 0.908 | 0.354 | 0.262 |
| Elevated D-dimer | 0.697 | <0.001 | 0.851 | 0.542 | 0.393 |
| BMI (kg/m<sup>2</sup>) | 0.558 | <0.001 | 0.695 | 0.434 | 0.129 |

Based on the multivariable regression results, a prognostic nomogram was established by integrating independent risk factors, including multiple surgeries, varicose veins, Preoperative hospital stay, age, BMI, trauma surgery, general surgery, neurosurgery, polycythemia, thrombocytosis, increased FDP, elevated D-dimer, hypoxemia, and elevated IL-6. To estimate the probability of thrombosis for an individual patient, a vertical line is drawn upward from the value of each variable to determine its corresponding score on the Points axis. The individual scores are subsequently accumulated to calculate a Total Points score. Finally, the point on the risk axis aligned with the total score represents the specific probability of thrombosis risk.

The predictive performance of the constructed nomogram was comprehensively evaluated using receiver operating characteristic (ROC) curves, calibration curves, and decision curve analysis (DCA).

The ROC curve analysis yielded an area under the curve (*AUC*) of 0.856 (95% CI: 0.843–0.870), demonstrating favorable discriminative ability. At the optimal cut-off value of 0.123, the model exhibited a sensitivity of 0.873 and a specificity of 0.690. (Figure 1)

**Figure 1.**
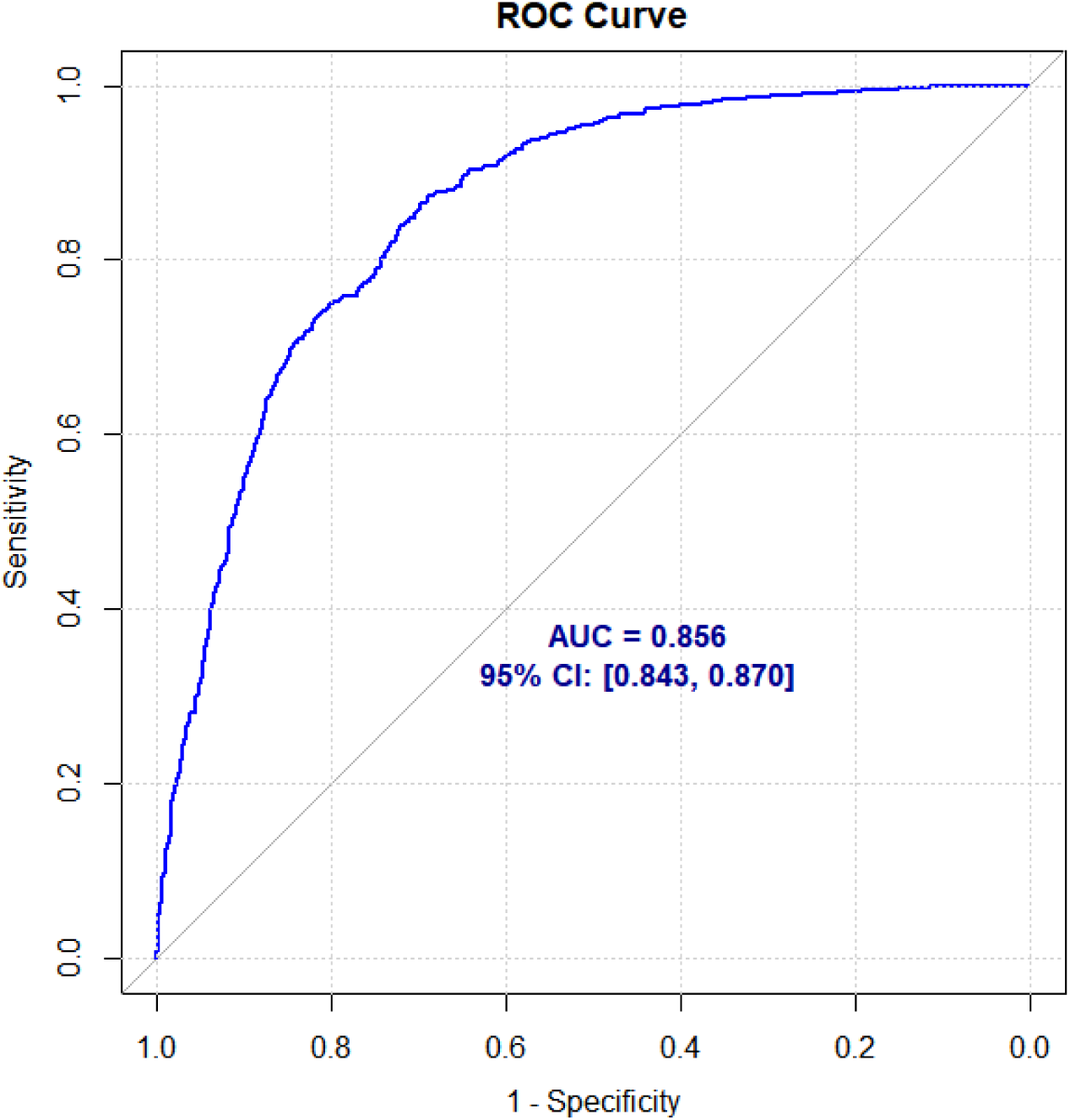

The calibration plot reflected the high agreement between the predicted risk of thrombosis and the true clinical observations. In the plot, the horizontal and vertical axes represent the predicted and observed probabilities, respectively. While a perfect calibration is represented by the 45° diagonal line(Figure 2), the bias-corrected curve showed negligible deviation from this ideal baseline, indicating excellent calibration accuracy. (Figure 4)

**Figure 2.**
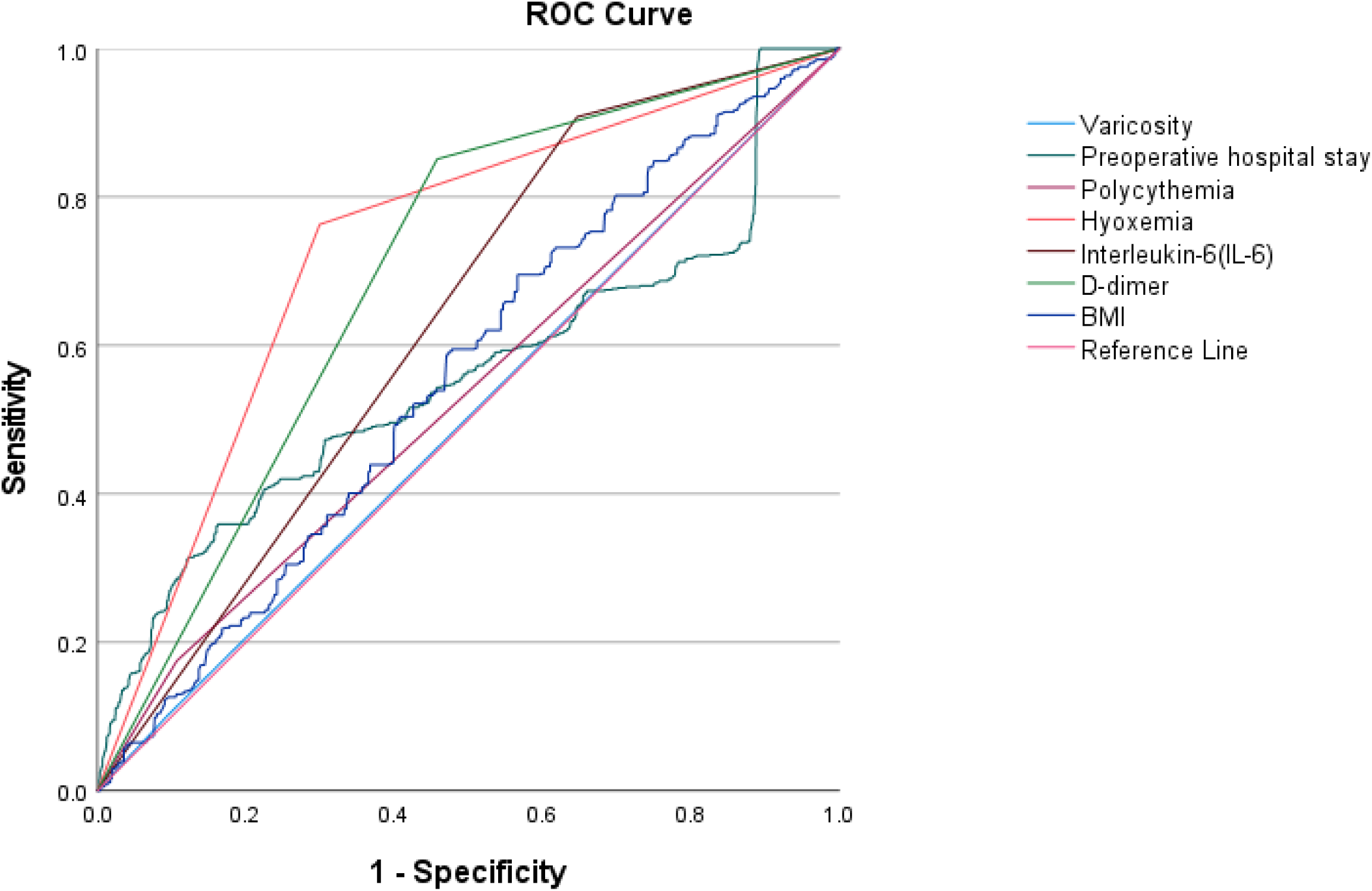

Based on the results of logistic regression analysis, an nomogram model was constructed using independent predictors. In the figure, each category of a variable corresponds to a point on the respective axis of the nomogram; drawing a vertical line upward from this point yields the score for that variable. The total score is obtained by summing the individual scores of all variables. The corresponding point on the risk axis for this total score indicates the probability of thrombosis. (Figure 3)

**Figure 3.**
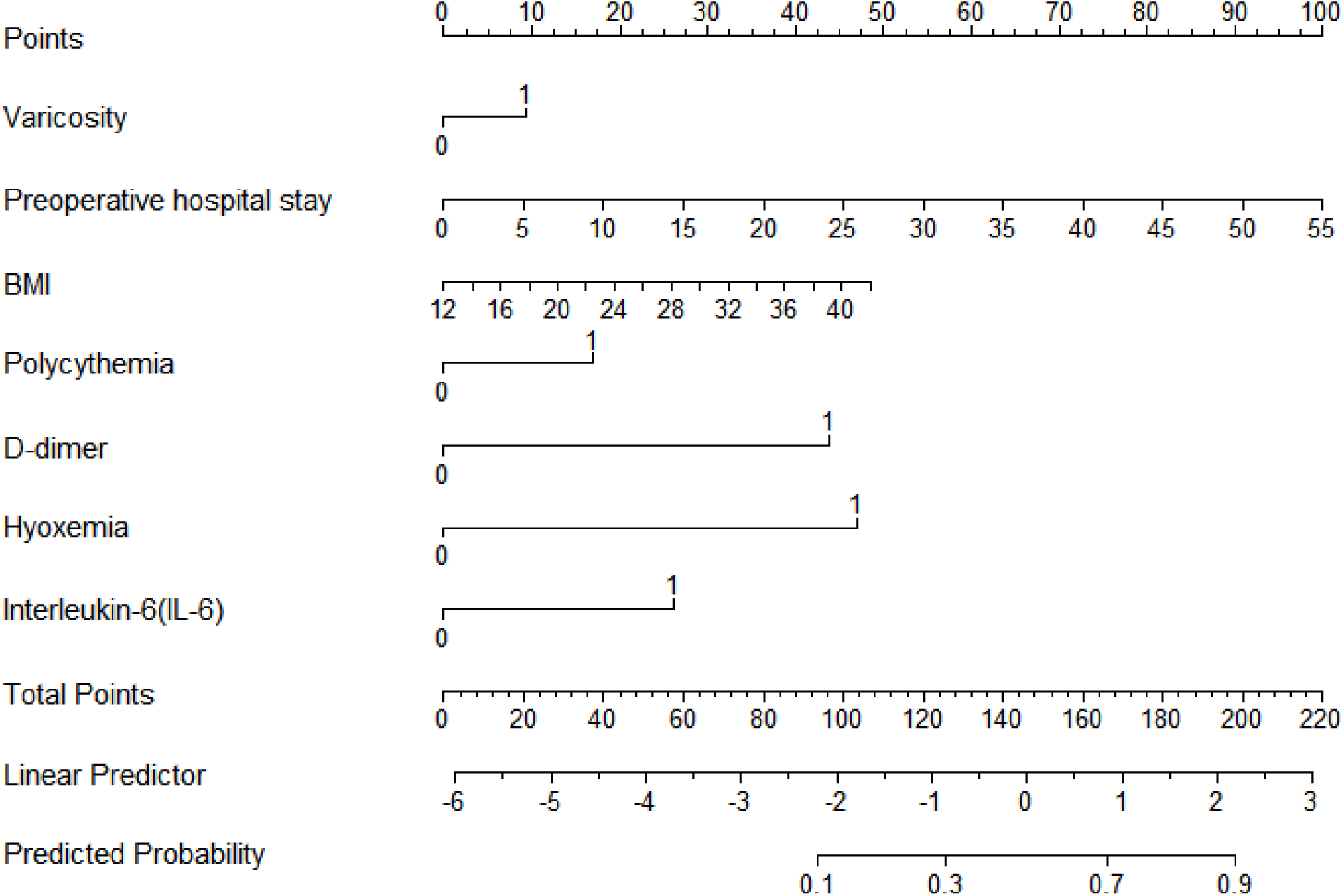

**Figure 4.**
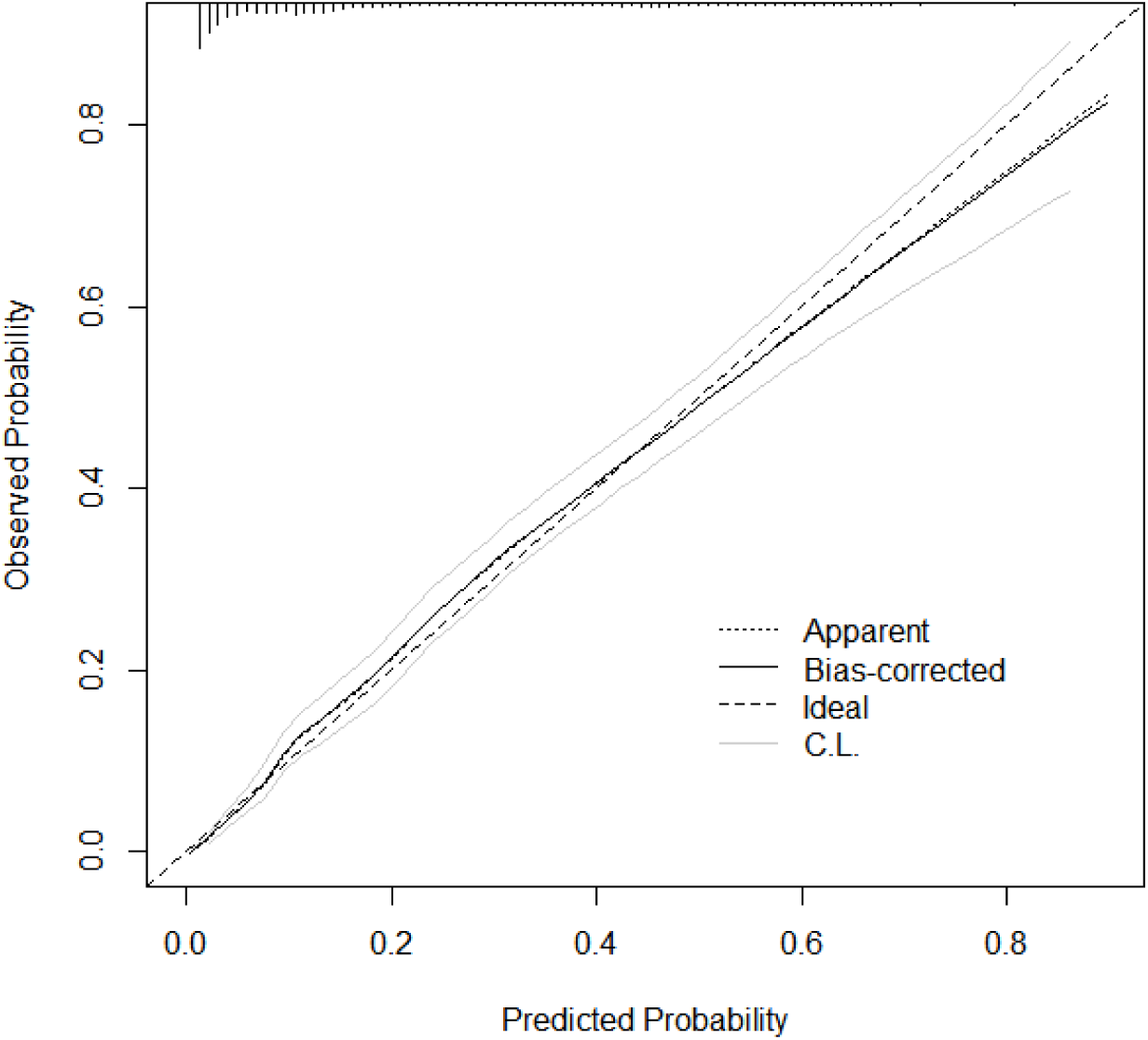

To evaluate clinical net usefulness, decision curve analysis (DCA) was employed to weigh potential benefits against intervention risks. In this analysis, the “None” baseline represents a default of zero interventions, whereas the “All” baseline assumes intervention for the entire cohort. As shown in the decision curve, the model outperformed both the “None” and “All” strategies, yielding a superior net benefit across the threshold probability interval from 0 to 1. (Figure 5)

**Figure 5.**
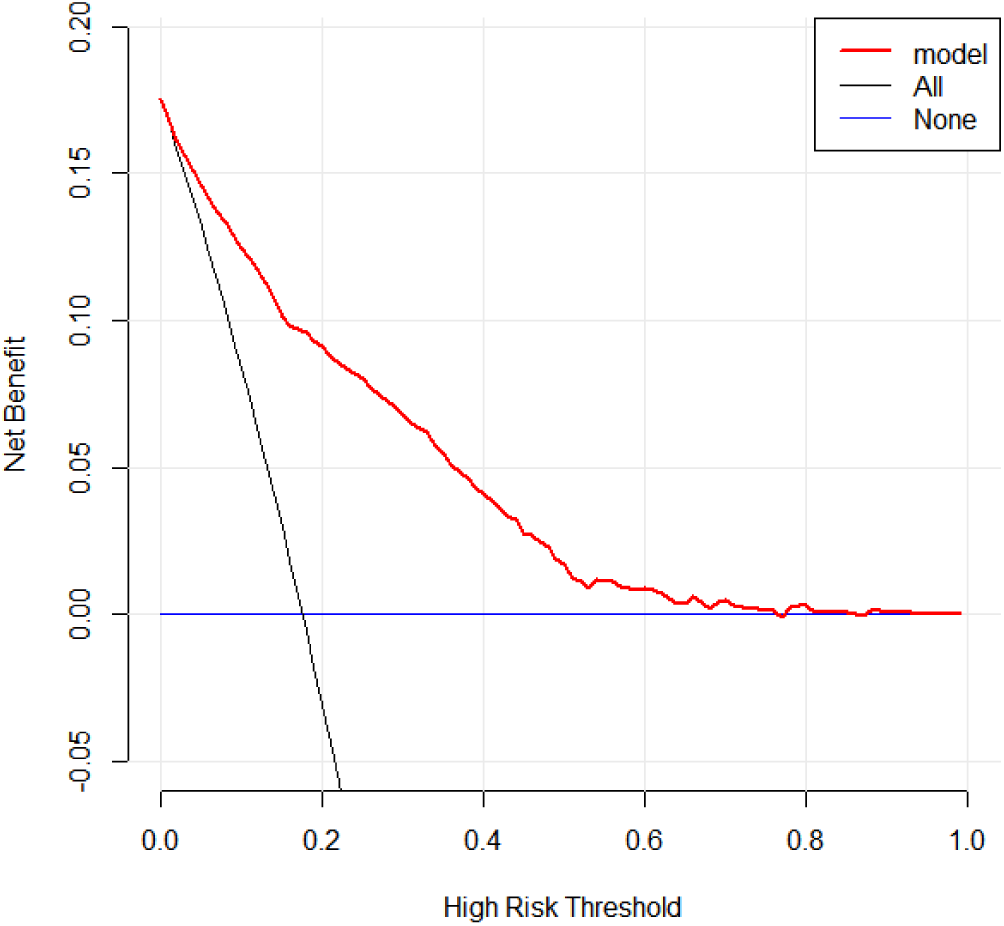

## Discussion

Perioperative venous thrombosis (VT) is a critical complication associated with significant mortality and morbidity [5–7]. While various risk assessment models are currently available, they are predominantly validated in low-altitude regions and omit high-altitude-specific variables, including residential altitude, hypoxemia, and polycythemia. In this context, promptly identifying high-risk patients and executing tailored interventions are essential to improving patient prognosis in high-altitude medical settings.

Considering the relatively low incidence of thrombosis across various subtypes, and acknowledging that any type of thrombosis presents a significant risk to patient safety, we conducted a combined analysis to ensure that the multivariate analysis possessed adequate statistical power. We retrospectively analyzed data from 11,084 patients who underwent surgical procedures. A multivariate logistic regression model was developed, incorporating residential altitude, hypoxemia, and other clinical variables.. The multivariable regression revealed that varicose veins, Preoperative hospital stay duration, BMI, polycythemia, elevated D-dimer, hypoxemia, and elevated IL-6 were independent risk factors for high-altitude perioperative thrombosis (*P* < 0.05). Among these, hypoxemia possessed the strongest predictive power, characterized by an OR of 7.249 (95% CI}: 5.936-8.852, *P* < 0.001) and the highest SHAP value contribution. Individual ROC curves indicated that hypoxemia yielded the highest diagnostic performance among single indicators, with an *AUC* of 0.732. Integrating these risk factors into a joint predictive model significantly upgraded the *AUC* to 0.856 (95% CI:0.843-0.870), demonstrating the superiority of a comprehensive risk assessment. Lastly, decision curve analysis (DCA) revealed that the model curve outperformed both the “all-intervention” and “no-intervention” strategies across the threshold probabilities, verifying its excellent clinical applicability.

Both hospitalization and surgery represent definitive risk factors for venous thromboembolism (VTE), with major surgical procedures further compounding this risk through prolonged immobilization, vascular endothelial damage, and acute hypercoagulable states[8–12]. At high altitudes, this baseline risk is uniquely amplified by the hypoxic and low-temperature climate, which directly instigates the elements of Virchow’s triad: hypercoagulability, endothelial injury, and stasis of blood flow [13–15]. Hypoxic exposure drives compensatory polycythemia and hemorheological alterations while impairing endothelial integrity[16], serving as a critical mechanism behind the increased perioperative thrombotic risk. Over time, high-altitude populations develop a series of secondary conditions, such as pulmonary hypertension, chronic cor pulmonale, and high-altitude polycythemia[17, 18]. These clinical entities engage in a complex interaction and cascade effect with chronic hypoxia[19].

From a mechanistic perspective, hypoxia-driven molecular alterations create a receptive milieu for inflammation and coagulation, largely mediated by the interaction between the HIF and NF-κB pathways[20]. Under hypoxic conditions, the increased endothelial expression of ICAM-1 and VCAM-1 enhances the adherence of leukocytes and platelets to the vessel wall, exacerbating thrombus formation[21]. Simultaneously, the inhibition of nitric oxide (NO) production impairs endothelial vasodilation, resulting in reduced blood flow velocity [22, 23]. Recent studies highlight that HIF-1α triggers NLRP3 inflammasome activation, which accelerates

TF/FVII expression via the transcription factor Egr-1, reinforcing platelet activation and driving the clotting cascade[24]. Additionally, HIF-1α-mediated upregulation of plasminogen activator inhibitor-1 (PAI-1) halts fibrinolytic activity, contributing to the structural stability of the formed thrombus[25] ^[25]^. Hypoxia also stimulates platelet autophagy, locking platelets into a hypercoagulable phenotype that facilitates aggregation [26]. Specifically, elevated blood viscosity and slowed flow velocity secondary to compensatory polycythemia[27–29], these pathways converge to create optimal conditions for thrombosis.

It is worth noting that while oxygen therapy is a mainstay of clinical management during hospitalization, the alternating exposure to hypoxia and hyperoxia may provoke ischemia-reperfusion injury (IRI). This process severely worsens endothelial dysfunction[30], thereby elevating the risk of thrombus formation[31, 32] IRI-induced reactive oxygen species (ROS) bursts damage endothelial integrity and trigger a transition to a procoagulant phenotype. This molecular shift promotes platelet activation and aggregation, activates both inflammatory responses and the clotting cascade, and concurrently upregulates PAI-1 to impair fibrinolytic function[26, 33]. When compounded by the exacerbation of hypoxemia caused by surgery and anesthesia, the risk of thrombosis accumulates substantially. Ultimately, this study is the first to establish an independent correlation between hypoxemia and thrombosis in a high-altitude setting, revealing the pivotal clinical value of recognizing hypoxia as a key risk factor.

Beyond hypoxemia, our multivariable analysis confirmed that varicose veins, length of hospital stay, polycythemia, BMI, elevated IL-6, and elevated D-dimer were independent risk factors for perioperative thrombosis. **(1)** Varicose veins, a typical manifestation of chronic venous disease, are closely linked to blood stasis and endothelial impairment [34]. In a high-altitude hypoxic climate, venous blood flow slows down further and red blood cell aggregation increases, significantly raising the risk of thrombosis in these patients. Here, the *OR* for varicose veins was 1.483, 95%CI: 1.075-2.047;, suggesting that clinicians should place greater emphasis on early screening and management for this population. **(2)** The positive correlation between the preoperative hospital stay duration and thrombosis (*OR* = 1.078, *P* < 0.001) reflects the cumulative impact of iatrogenic immobilization on the coagulation cascade. At high altitudes, the delayed recovery of postoperative pulmonary function and diminished exercise tolerance frequently prolong hospitalization, further compounding the baseline thrombotic risk. This indicates that optimizing perioperative care pathways and promoting early mobilization are exceptionally paramount in high-altitude medical practice. **(3)** Although polycythemia is a common compensatory change among high-altitude residents, excessive polycythemia causes a marked increase in blood viscosity and resistance, thus driving clot formation[35, 36]. Our study confirmed polycythemia as an independent risk factor (*OR* = 2.097, *P* < 0.001), which fits perfectly with the pathophysiological process where chronic hypoxia stimulates red blood cell production. **(4)** While the association between a higher BMI and thrombosis is well-documented in low-altitude areas, obese patients at high altitudes often present with sleep apnea syndrome and daytime hypoxemia.

The interaction between hypoxia and obesity can further expand the risk of thrombosis[37]. In our model, each unit increase in BMI was associated with a 7.2% increase in the odds of thrombosis (*OR* = 1.072), showing that weight control is also vital during the perioperative period at high altitudes. **(5)** IL-6 is a key driver of inflammation, and surgical trauma can induce its massive release, which in turn sets off the coagulation cascade[38]. In this study, elevated IL-6 increased the risk of thrombosis by 2.964 times (*OR* = 2.964, *P* < 0.001). Because hypoxia itself upregulates inflammatory factors at high altitudes [39], it overlaps with surgery-induced stress to create a dangerous “hypoxia-inflammation-thrombosis” cycle. **(6)** Elevated D-dimer levels, indicating fibrinolytic system activation, have been universally applied to screen for thrombosis. In our study, the OR for elevated D-dimer was 6.203 (95% CI: 4.852–7.932), demonstrating that D-dimer maintains excellent predictive performance in high-altitude medical settings.

Intriguingly, an unexpected finding of this study is that the altitude of residence did not remain in the final regression model and was not identified as an independent risk factor for perioperative thrombosis. To guarantee the reliability of this observation, we adopted two different modeling strategies for the altitude variable in our univariate screening. We first examined residential altitude as a continuous variable to test for a linear trend; we then categorized patients into “high-altitude” (1,500–2,500 meters) and “plateau” (>2,500 meters) subgroups, evaluating it as a dichotomous variable (0/1). In both instances, no statistically significant correlation with perioperative thrombosis was observed (*P* > 0.05). Although this result seems to contradict the long-held paradigm that living at higher altitudes escalates thrombotic risk, it aligns perfectly with our finding that hypoxemia is the independent risk factor. In essence, the pathogenic impact of geographical altitude is entirely mediated by the resulting drop in PaO_2_; once PaO_2_ is accounted for, the direct effect of altitude disappears. Beyond this biological mediation, several multifaceted factors may account for why geographical altitude per se did not emerge as an independent predictor in the final model:

First, alterations in environmental exposure introduce a major confounding effect. Patients frequently experience a significant shift in their physical surroundings when traveling from their diverse residential locations to the clinical site. Given that the hospital is located at an elevation of roughly 2,200 meters, every patient is exposed to the same standardized hypoxic baseline upon admission. For patients accustomed to much higher elevations, hospitalization actually brings about a relative rise in barometric oxygen pressure. On the other hand, patients arriving from lower altitudes undergo a relative decrease in oxygen exposure. This dynamic environmental transition means that a patient’s home altitude cannot reliably serve as a surrogate for their actual perioperative hypoxic exposure level.

Second, there is a fundamental distinction between chronic adaptation and acute physiological stress. Individuals residing at high altitudes for extended periods generally develop robust physiological adaptations, such as compensatory polycythemia, increased minute ventilation, and heightened tissue oxygen extraction capacity. These long-term adjustments can partially counteract or offset the prothrombotic effects of environmental changes encountered during hospital admission. Ultimately, acute hypoxic stress during the perioperative period (triggered by anesthesia, surgical tissue injury, postoperative pain, and prolonged bed rest) provides a more direct and reliable prediction of thrombotic susceptibility than chronic geographical exposure alone.

Third, there is a distinct non-linear relationship between ambient altitude and hypoxemia. The association between where a patient lives and their true arterial oxygen tension (PaO_2_) is rarely a straightforward linear one, as it is confounded by multiple individual factors such as lung function, hemoglobin levels, and hypoxic ventilatory responses. Due to these diverse adaptive capabilities, some patients from extreme high-altitude areas manage to maintain well-compensated PaO_2_ levels upon hospital admission, showing no significant difference compared to patients from lower elevations. Ultimately, directly measuring PaO_2_ provides a far more reliable and clinically actionable reflection of a patient’s hypoxic status than a simple estimation based on geographical residence.

Fourth, limitations in the sample’s altitude composition may have obscured the true effect. The residential altitude of the majority of patients (roughly 85%) in this cohort was heavily concentrated between 1,500 and 3,000 meters. This narrow distribution range means there was insufficient variability in the altitude data to successfully reveal a statistically significant link to thrombus formation. Future studies recruiting a broader spectrum of individuals, particularly those from very low altitudes (< 500 meters) or extreme elevations (> 4,000 meters) are warranted to more clearly delineate the independent impact of geographical altitude on thrombotic risk.

## Limitations

Several limitations should be acknowledged when interpreting our findings. (1) The retrospective study design inherently limits the ability to obtain several established risk factors for thrombosis, such as a history of venous thromboembolism, thrombophilia, smoking status, and hormone therapy usage, from the study’s database. Consequently, these factors were excluded from the analysis, potentially introducing selection bias and information bias.(2) We utilized standard sea-level criteria to define hypoxemia; however, these cut-offs might not be perfectly tailored to residents well-adapted to high altitudes, potentially leading to an underestimation of hypoxemia’s actual burden. (3) This study did not undergo external validation, and the data were drawn from a single medical center. Therefore, further multi-center research is warranted in the future to confirm the robustness and generalizability of our predictive model.

## Summary

In summary, the development of perioperative thrombosis at high altitudes is a complex interplay of multiple risk factors, driven fundamentally by hypoxemia. Other critical elements, such as varicose veins, length of stay, polycythemia, BMI, elevated IL-6 levels, and elevated D-dimer, interface and converge through diverse pathophysiological routes. To mitigate this risk, clinical prevention efforts should center on the core objective of reversing hypoxemia, alongside tailored, patient-specific control of these co-existing clinical risk factors.

